# Hyperactivity of mTORC1 and mTORC2-dependent signaling mediate epilepsy downstream of somatic PTEN loss

**DOI:** 10.1101/2023.08.18.553856

**Authors:** Erin R. Cullen, Mona Safari, Isabelle Mittelstadt, Matthew C. Weston

## Abstract

Gene variants that hyperactivate PI3K-mTOR signaling in the brain lead to epilepsy and cortical malformations in humans. Some gene variants associated with these pathologies only hyperactivate mTORC1, but others, such as *PTEN*, *PIK3CA*, and *AKT*, hyperactivate both mTORC1- and mTORC2-dependent signaling. Previous work established a key role for mTORC1 hyperactivity in mTORopathies, however, whether mTORC2 hyperactivity contributes is not clear. To test this, we inactivated mTORC1 and/or mTORC2 downstream of early *Pten* deletion in a new model of somatic *Pten* loss-of-function (LOF) in the cortex and hippocampus. Spontaneous seizures and epileptiform activity persisted despite mTORC1 or mTORC2 inactivation alone, but inactivating both mTORC1 and mTORC2 simultaneously normalized brain activity. These results suggest that hyperactivity of both mTORC1 and mTORC2 can cause epilepsy, and that targeted therapies should aim to reduce activity of both complexes.

## Introduction

Genetic disruptions that hyperactivate mTOR activity, termed ‘mTORopathies’, cause macrocephaly, intractable childhood epilepsies, and behavioral presentations including autism spectrum disorders [1–4]. The mTOR kinase can function in two complexes, mTORC1 and mTORC2, each with unique downstream targets and upstream activators [5]. A unifying feature of mTORopathies is hyperactivation of mTORC1, which has led to the hypothesis that this complex and its effectors mediate disease phenotypes [6]. A subset of mTORopathies however, (e.g. those caused by variants in *PTEN*, *PIK3CA*, and *AKT*), hyperactivate both mTORC1 and mTORC2 [7,8], and mTORC2 hyperactivity has also been reported in human TLE [9]. It is unclear whether mTORC2 hyperactivity contributes to, or can cause, disease phenotypes such as epilepsy.

*Pten* LOF in neurons induces soma hypertrophy, increased dendritic branching, and synaptic hyperconnectivity [10,11]. These phenotypes resemble those observed in models of mTORC1-specific hyperactivation [12], and can be prevented with either the mTORC1 inhibitor rapamycin or loss of the mTORC1-specific protein RAPTOR [13]. Rapamycin also rescues epilepsy and mortality associated with *Pten* loss, even after epilepsy has been established, suggesting that mTORC1 hyperactivity underlies these phenotypes [14–16]. Thus, there is strong evidence that hyperactivation of mTORC1 downstream of PTEN disruption causes the macrocephaly, epilepsy, early mortality, and synaptic dysregulation observed in humans and model organisms [17].

There is also evidence against the mTORC1-centric hypothesis. Rapamycin may affect mTORC2 activity at high doses or over long periods of time [18,17], and mTORC2 regulates synapse function, neuronal size, and cytoskeletal organization independent of mTORC1 [19,20]. In a mouse model in which *Pten* was inactivated in forebrain neurons in early adulthood, mTORC2 inhibition, but not mTORC1 inhibition, rescued seizures and behavioral abnormalities. In this model, epilepsy was dissociated from macrocephaly, which was rescued by mTORC1 inhibition but not mTORC2 inhibition [21]. However, mTORC2 inactivation did not normalize spontaneous seizures or seizure susceptibility in a model of *Pten* loss in dentate granule neurons [22]. Thus, there is conflicting evidence about whether mTORC1 or mTORC2 hyperactivity causes epilepsy downstream of *Pten* loss.

To address this, we suppressed mTORC1 and mTORC2 activity alongside *Pten* loss by inducing simultaneous deletion of *Rptor* or *Rictor*, whose protein products are unique and essential components of mTORC1 and mTORC2, respectively, in a model of somatic *Pten* LOF. In this model, epilepsy could be rescued by concurrent mTORC1 and mTORC2 inactivation, but persisted when either gene remained intact, suggesting that hyperactivity of either complex can lead to neuronal hyperexcitability and epilepsy.

## Results

*Generation of a developmental brain somatic mosaic model of Pten LOF.* To model the epileptogenic somatic mutations often observed in mTORopathies [23,24], we injected an AAV9 virus expressing GFP and Cre under control of the *hSyn* promoter into one hemisphere of the cortex of *Pten^(fl/fl)^, Pten^(fl/fl)^-Rptor^(fl/fl)^, Pten^(fl/fl)^-Rictor^(fl/fl)^,* and *Pten^(fl/fl)^-Rptor^(fl/fl)^-Rictor^(fl/fl)^*mice at P0 (Fig. 1A). We hereafter refer to these mice as Pten LOF, Pten-Rap LOF, Pten-Ric LOF, and PtRapRic LOF. Green fluorescent protein (GFP) was expressed in neurons throughout the cortical layers in all experimental animals, but largely confined to one cortical hemisphere and the underlying hippocampus (Fig. 1B and Supplementary Fig. 1). The percentage of cells that were GFP+ was similar across groups, although there was a decrease in cell density in the Pten LOF and Pten-Ric LOF groups [25] (Fig. 1C).

**Figure 1.**
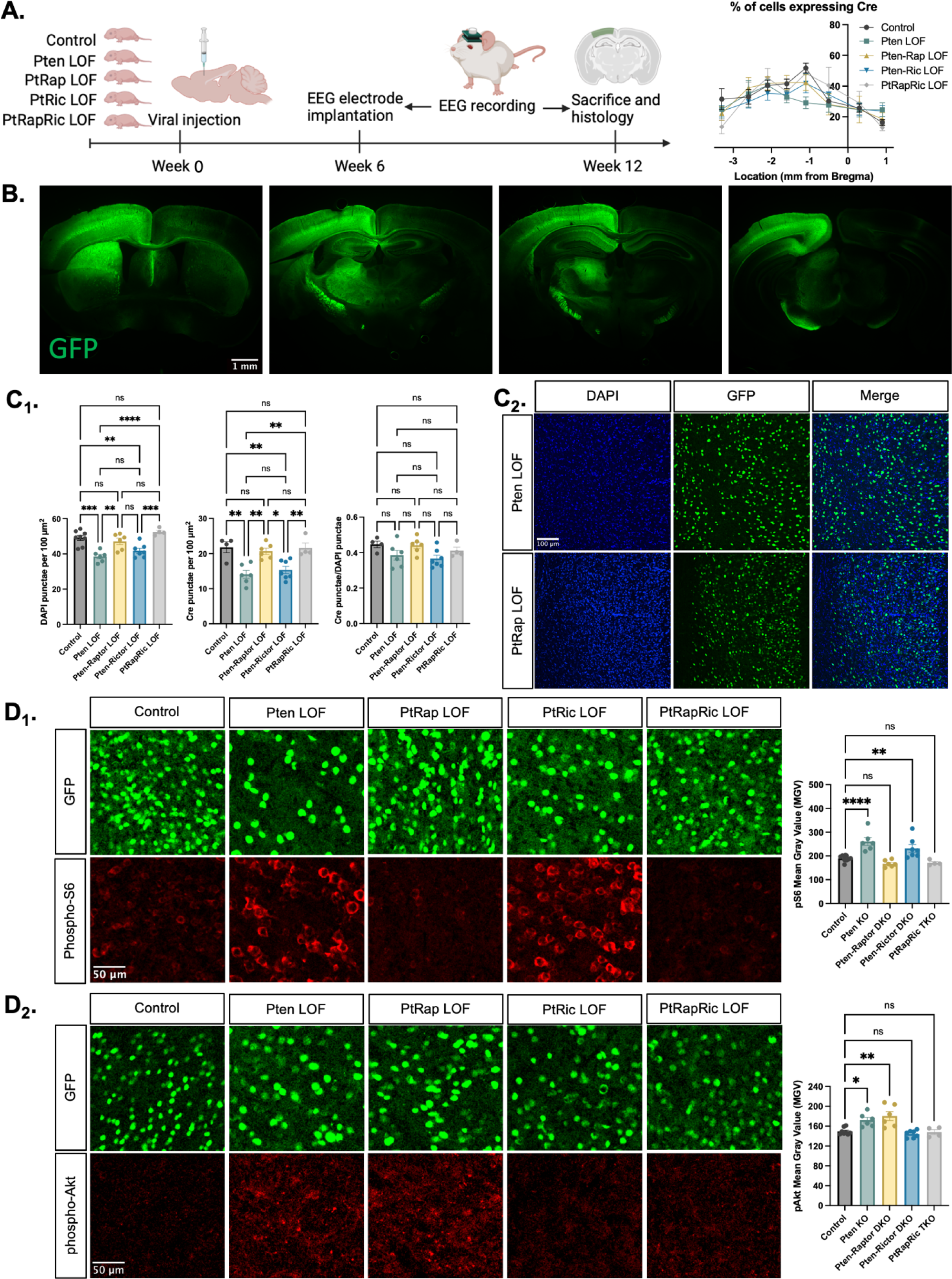
Histological characterization of the focal cortical Pten LOF model. A) Experimental timeline showing induction of hSyn-Cre-GFP or Control hSyn-GFP AAV at P0 and EEG recording in adulthood. The location of cortical Cre expression relative to Bregma did not differ between groups. B) Representative images of GFP expression in a Control mouse brain, demonstrating expression predominantly in one hemisphere of the cortex. Projections from affected neurons can be seen in white matter tracts including the fornix, internal capsule, corpus callosum, and cerebral peduncle. C1) Quantification of lesion severity and neuron density in Cre-expressing animals. There were fewer Cre-expressing neurons per unit area in the Pten LOF and Pten-Ric LOF groups, which was at least partially attributable to a decrease in cell density in these groups. No significant differences in Cre expression remained when Cre expression was calculated based on cell density rather than area. C2) Representative images of DAPI and Cre fluorescence in the cortex of Pten LOF and Pten-Rap LOF animals. Cell density and Cre density in Pten-Rap LOF animals was indistinguishable from Controls. D1) Phospho-S6, a marker of mTORC1 activity, was increased by Pten LOF and reduced to control levels by concurrent *Rptor* loss. Phospho-S6 was also increased from Control levels in Pten-Ric LOF, indicating that mTORC1 hyperactivity was not normalized by *Rictor* loss. Combined *Rptor/Rictor* loss also normalized phospho-S6 expression. D2) phospho-Akt, a marker of mTORC2 activity, was increased in Pten LOF and normalized by *Rictor* loss, but not by *Rptor*. Combined *Rptor/Rictor* loss also normalized phospho-Akt expression. Error bars show mean ± s.e.m. ns indicates p>0.05, * indicates p<0.05, ** indicates p<0.01, *** indicates p<0.001, and **** indicates p<0.0001 as assessed by statistical tests indicated in Table 3. Diagram created with BioRender.com.

We analyzed the impact of *Pten*, *Rptor*, and *Rictor* LOF on mTORC1 and mTORC2 activity in the affected hemisphere using quantitative immunohistochemistry. Phospho-S6 levels, which report mTORC1 activity through S6K1, were increased from Controls in Pten LOF and Pten-Ric LOF brains, but rescued in Pten-Rap LOF brains (Fig. 1D_1_). Phospho-Akt (S473) levels, which report mTORC2 activity, were increased in Pten LOF and Pten-Rap LOF but normalized in Pten-Ric LOF brains (Fig. 1D_2_). Levels of both pS6 and pAkt were not different from Controls in PtRapRic LOF brains. These results indicate that *Pten* LOF hyperactivates both mTORC1 and mTORC2 in this model, and that these increases in activity are independently rescued by mTORC1 and mTORC2 inactivation.

*mTORC1 and/or mTORC2 inactivation attenuate Pten LOF-induced macrocephaly, cellular overgrowth, and electrophysiological changes.* Macrocephaly and neuronal hypertrophy are well-characterized sequelae of deficient PTEN signaling in patients and mouse models [26,27]. In mouse models of *Pten* LOF, these phenotypes can be rescued by rapamycin treatment or *Rptor* LOF [21,28,17,13]. Here, cortical thickness was increased in Pten LOF mice but reduced to Control levels in Pten-Rap LOF, Pten-Ric LOF, and PtRapRic LOF mice (Fig. 2A-B). In individual neurons, Pten LOF showed a ≈100% increase in mean soma size versus Control. Pten-Ric LOF significantly reduced soma size from Pten LOF levels, but still showed a 60% increase over Controls. Soma size in Pten-Rap LOF and PtRapRic LOF neurons did not significantly differ from Controls (Fig. 2C). Whole cell current clamp analysis of neuronal membrane excitability of GFP+ neurons in acute brain slices supported these conclusions, as input resistance, capacitance, and rheobase were all significantly altered by Pten LOF, but not different from control levels in Pten-Rap LOF, Pten-Ric LOF and PtRapRic LOF (Figure 2D). Spontaneous EPSCs were also recorded with whole cell voltage clamp.

**Figure 2.**
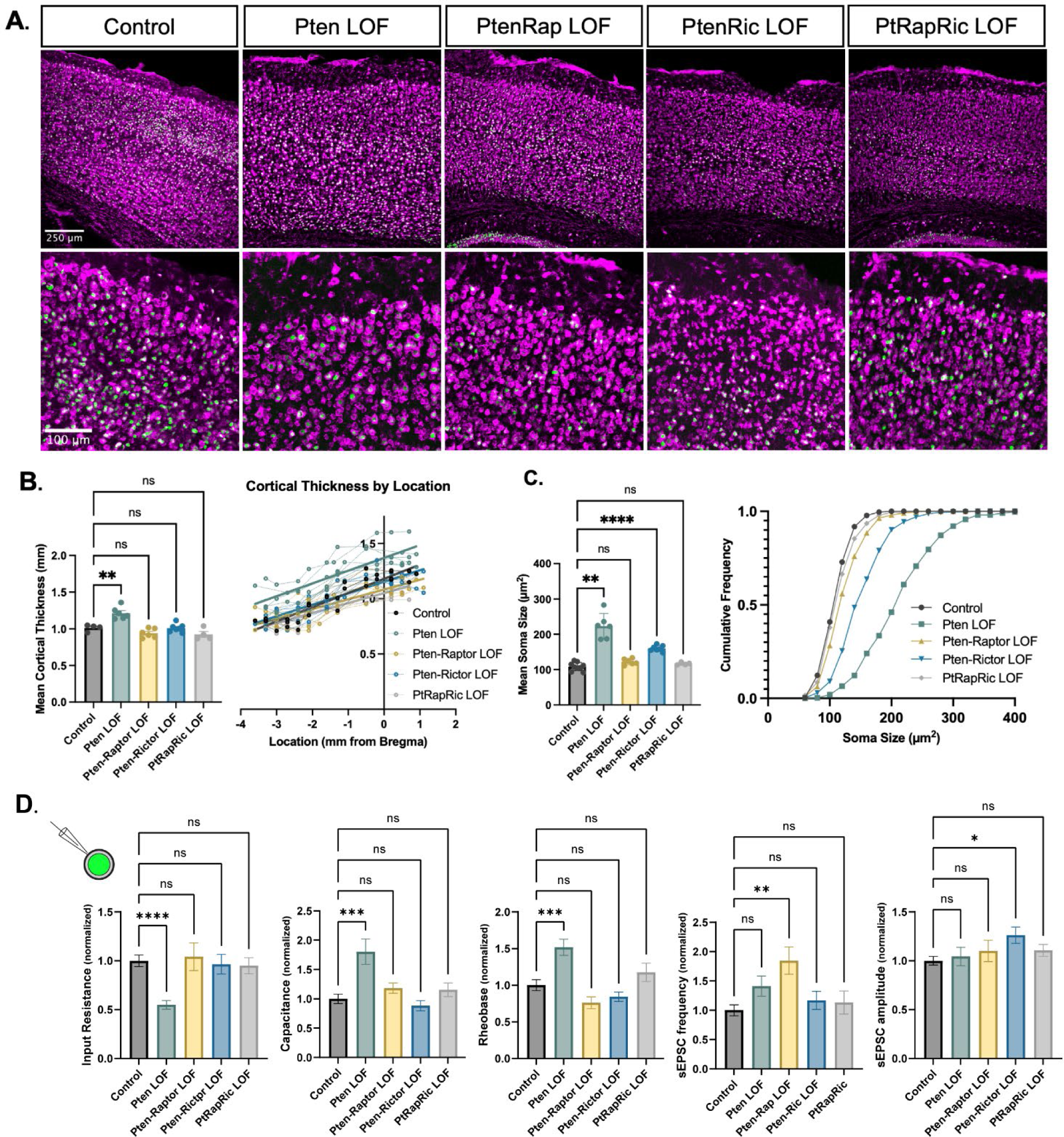
Independent mTORC1 or mTORC2 inactivation prevents most cellular effects of Pten LOF, but only dual mTORC1/2 inactivation prevents all. A) Example images showing fluorescent Nissl stain (magenta) and GFP expression (green) in cortical neurons. The top row shows the cortical thickness in all 5 groups and the bottom row shows zoomed in images depicting the differences in soma size across groups. B) The mean cortical thickness was increased in Pten LOF throughout the cortex. Pten-Rap, Pten-Ric, and PtRapRic LOF cortical thickness did not differ significantly from Controls. C) The mean soma size was strongly increased in Pten LOF and to a smaller extent in Pten-Ric LOF. Pten-Rap LOF and PtRapRic LOF groups did not differ significantly from Controls. D) Whole cell patch clamp analysis of GFP+ neurons showed that capacitance and rheobase were increased in Pten LOF neurons, whereas input resistance was decreased. These changes were not found in in Pten-Rap LOF, Pten-Ric LOF and PtRapRic LOF groups. The frequency of spontaneous EPSCs was elevated in Pten-Rap LOF neurons, and their amplitude was larger in Pten-Ric neurons. Patch data is normalized to values from littermate controls. Error bars show mean ± s.e.m. ns indicates p>0.05, * indicates p<0.05, ** indicates p<0.01, and **** indicates p<0.0001 as assessed by statistical tests indicated in Table 3.

Pten-Rap LOF and Pten-Ric LOF had elevated sEPSC frequency and amplitude, respectively, as previously reported [22,21,13]. Neither of these parameters were different from Control in PtRapRic LOF neurons (Figure 2D), indicating that only concurrent mTORC1/mTORC2 inactivation can normalize sEPSC parameters.

*Concurrent mTORC1/2 inactivation, but neither alone, prevents epilepsy and interictal EEG abnormalities in focal Pten LOF.* Next, we measured epileptic brain activity with video-EEG to determine the ability of mTORC1 or mTORC2 activity to rescue this key feature of mTORopathies, Generalized seizures (GS) were observed in 0/9 Control, 4/7 Pten LOF, 2/6 Pten-Rap LOF, 2/7 Pten- Ric LOF, and 0/6 PtRapRic LOF animals, suggesting that only simultaneous mTORC1 and mTORC2 inactivation potentially prevents GS. The frequency of GS events in Pten-Rap LOF and Pten-Ric LOF mice was not significantly lower than the GS frequency in Pten LOF mice, although the number of mice used was not powered to detect reductions in GS frequency. GS events were longer-lasting in Pten-Rap LOF animals than in Pten LOF or Pten-Ric LOF animals (Fig. 3A), and did not appear to be correlated with mTOR pathway activity (Supplementary Fig. 2).

**Figure 3.**
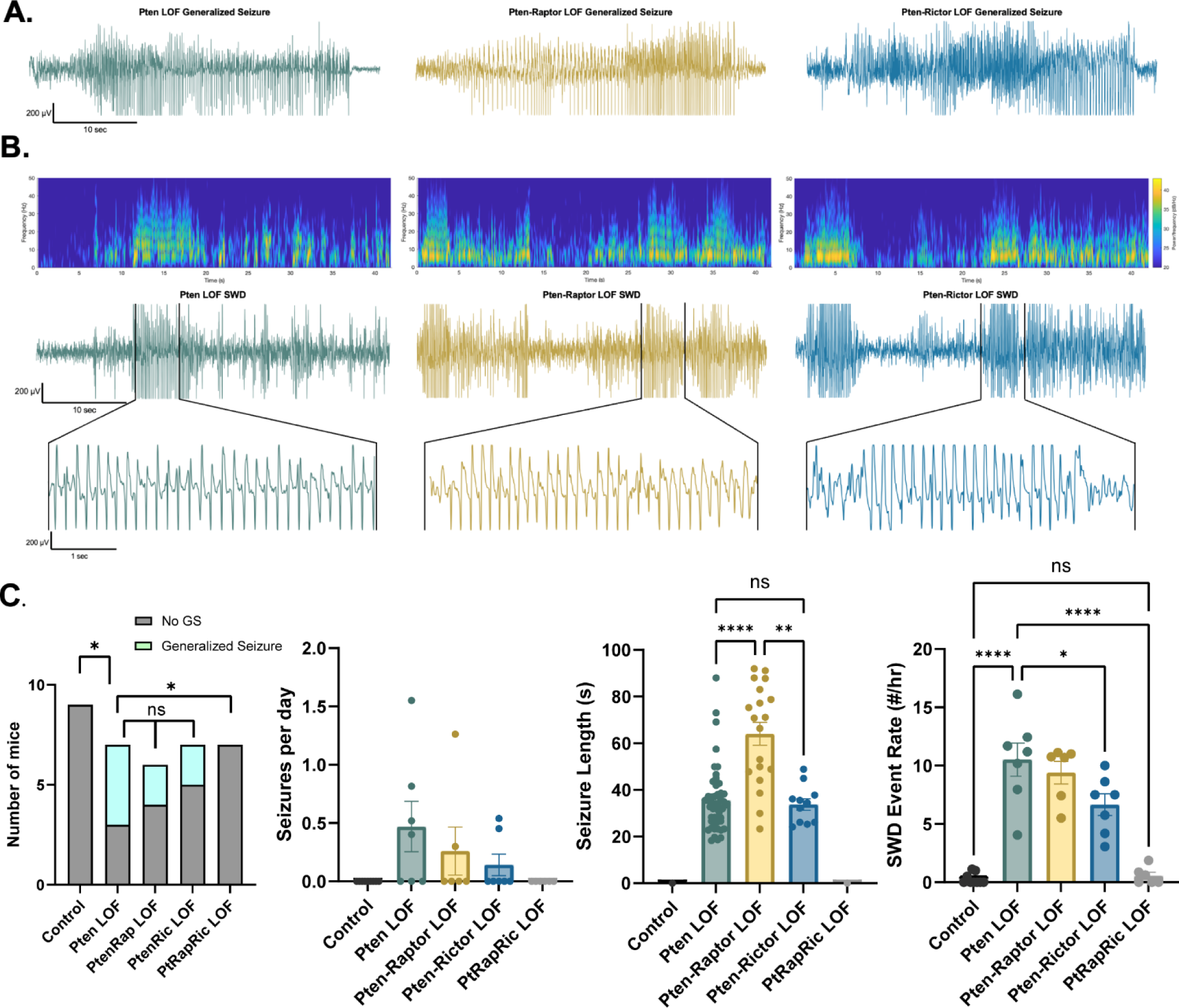
Combined mTORC1 and mTORC2 inactivation, but neither alone, rescues epilepsy in the focal Pten LOF model. Spontaneous seizures and interictal spike activity were assessed in Pten LOF, Pten-Rap LOF, Pten-Ric LOF, PtRapRic LOF, and Control mice. A) Representative traces of generalized seizures (GS) in a subset of animals in Pten LOF, Pten-Rap LOF, and Pten-Ric LOF groups. B) Spectrograms (top) and traces depicting example SWD event trains. C) Summary data showing the occurrence of GS per animal, number of GS per day, and GS length. GS events were significantly longer in the Pten-Rap LOF group than the other groups. In the Pten LOF group, 19% of GS events (9/48) exceeded 45 seconds in length, and these events were observed in 2/4 Pten LOF GS+ animals. 79% of GS events (15/19) in the Pten-Rap LOF group exceeded 45 seconds, and these events were observed in 2/2 GS+ Pten-Rap LOF animals. 1/11 GS events in the Pten- Ric LOF group exceeded this threshold. D. Summary data showing the SWD rate in all animals. Error bars show mean ±s.e.m. ns indicates p>0.05, * indicates p<0.05, ** indicates p<0.01, *** indicates p<0.001, and **** indicates p<0.0001 as assessed by tests indicated in Table 3.

The most striking and consistent type of epileptic brain activity in the Pten LOF animals was frequent 5-7 Hz spike trains lasting 3 seconds or more, which fit previous characterizations of spike-and-wave discharges (SWDs) in rodents [29]. SWDs were observed in all Pten LOF, Pten-Rap LOF, and Pten-Ric LOF animals, and in 3/9 Control and 3/6 PtRapRic LOF animals. The frequency of SWDs in Pten LOF, Pten-Rap LOF, and Pten-Ric LOF animals was significantly higher than Controls, but PtRapRic LOF completely blocked this increase. Pten-Ric LOF animals had a lower frequency of SWD events than Pten LOF animals (Fig. 3B). Taken together, these data indicate that inactivating mTORC1 provides no protection against *Pten* loss-induced epilepsy, and may even exacerbate it. mTORC2 inactivation provides some protection, but only concurrent mTORC1 and mTORC2 inactivation can prevent it.

In addition to epileptic activity, we also found that Pten LOF caused obvious alterations in features of the interictal EEG. To quantify these changes, we measured EEG coastline, absolute mean amplitude, and power spectra of the EEG. Coastline, mean amplitude, and total power were all significantly increased by Pten LOF, and these changes were not significantly reduced by Pten-Rap or Pten-Ric LOF. PtRapRic LOF, however, reduced these increases to Control levels. Correspondingly, EEG power was increased in Pten LOF animals (Fig. 4A; Table 1A). This increase was normalized by PtRapRic LOF, but not by Pten-Rap LOF or Pten-Ric LOF. PtRapRic LOF did not fully rescue a rightward shift in normalized EEG band power (Fig. 4B; Table 1B).

**Figure 4.**
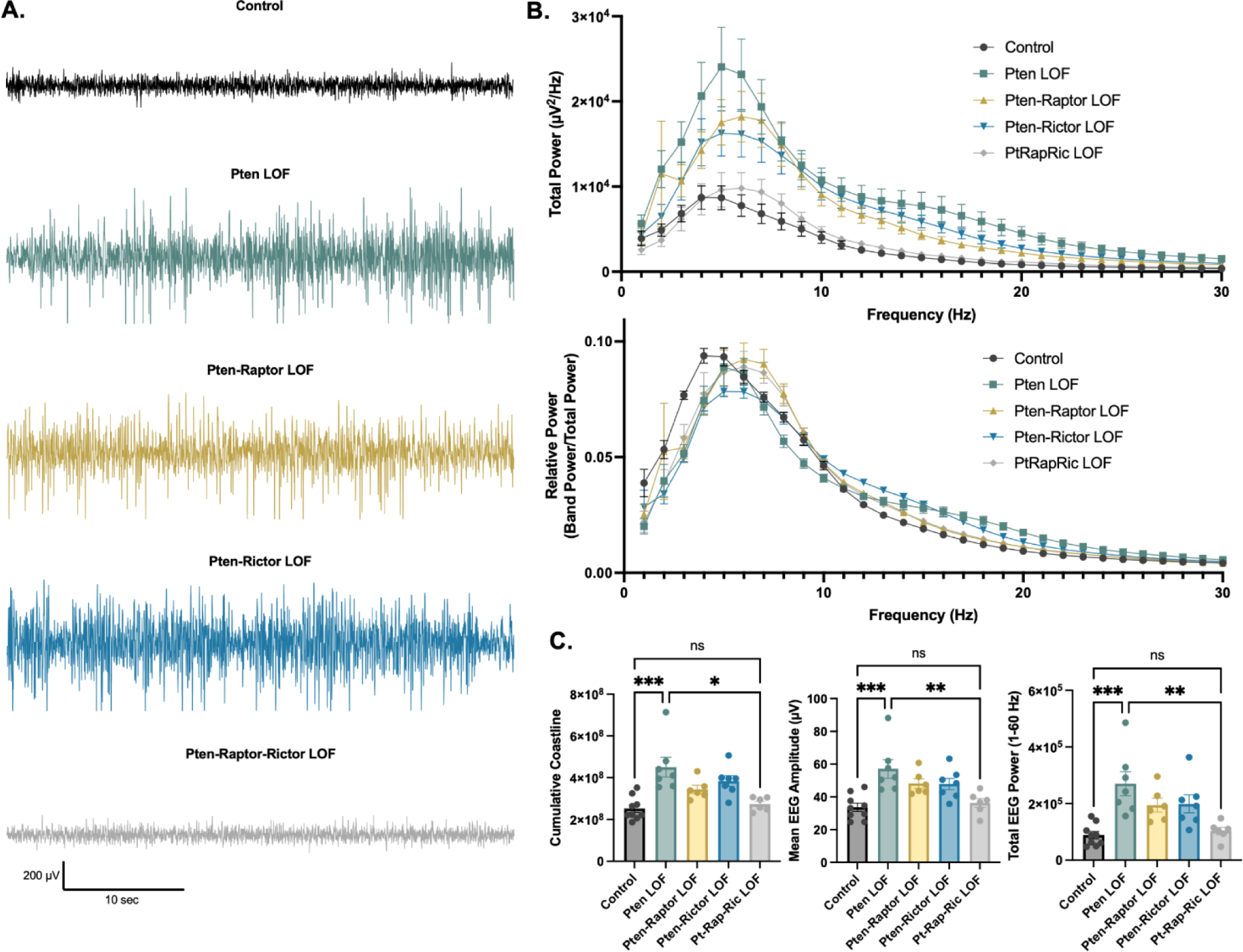
Combined mTORC1 and mTORC2 inactivation, but neither alone, rescues Pten LOF-induced abnormalities in the interictal EEG. A) Examples of typical EEG traces for each genotype. In EEG epochs that were not characterized as GTCS or SWD events, Pten LOF animals had higher levels of EEG activity as quantified by EEG line length, absolute mean amplitude, and total power. These changes were not significantly decreased in Pten-Rap LOF or Pten-Ric LOF mice, but were normalized in PtRapRic LOF mice. B) Total EEG power was increased by Pten LOF and attenuated, but not normalized, in either Pten-Rap or Pten-Ric LOF mice. Relative power was decreased in delta and increased in higher frequencies by Pten LOF. Pten-Rap LOF, Pten-Ric LOF, and PtRapRic LOF animals all showed a milder rightward shift of EEG power. C) Line length, mean amplitude, and power are increased in Pten LOF and normalized by PtRapRic LOF. Error bars show mean ± s.e.m. ns indicates p>0.05, * indicates p<0.05, ** indicates p<0.01, *** indicates p<0.001 as assessed by statistical tests indicated in Table 3. Two-way ANOVA p-values for EEG power are reported in Table 2.

**Table 1.**
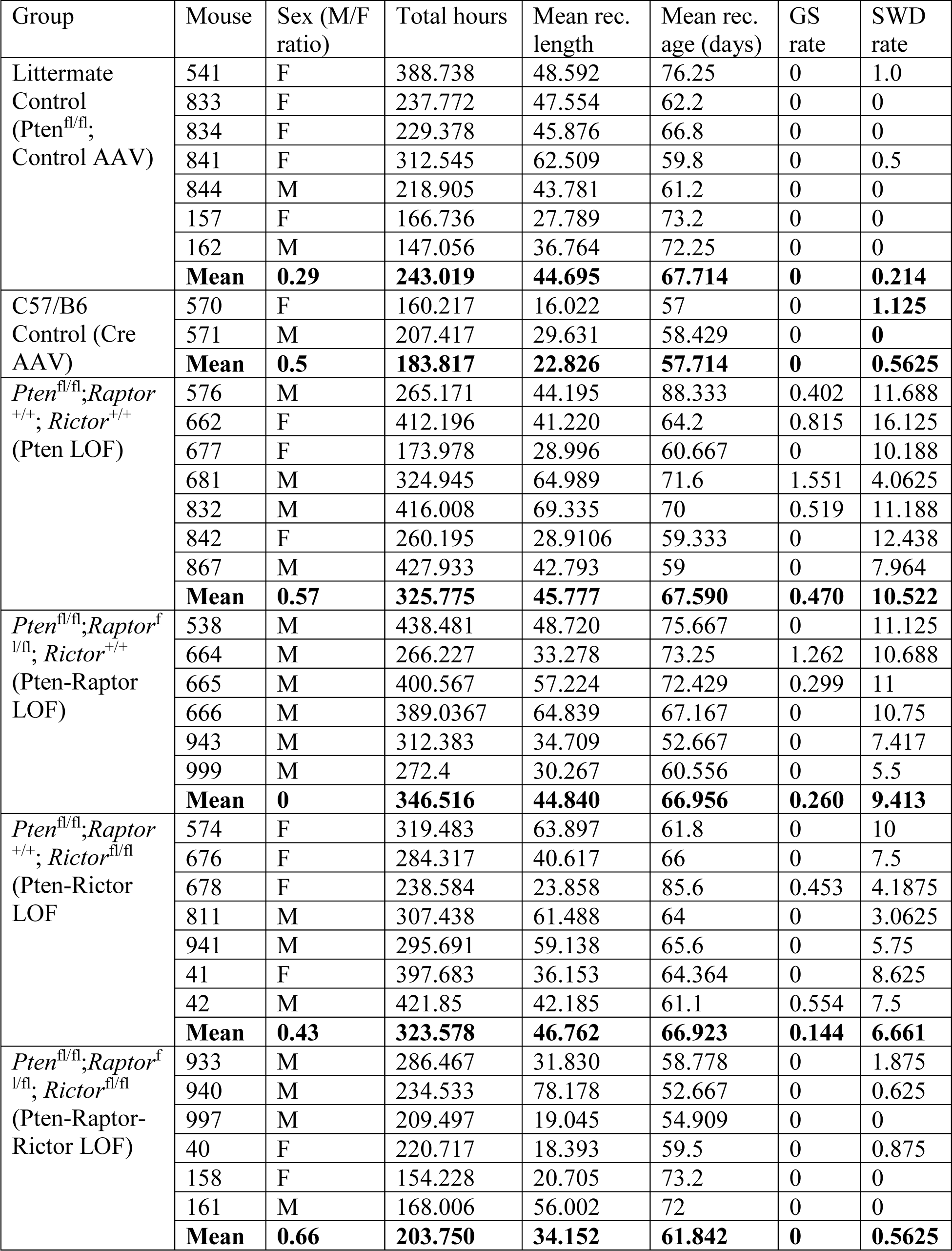
EEG Monitoring.

**Table 2.**
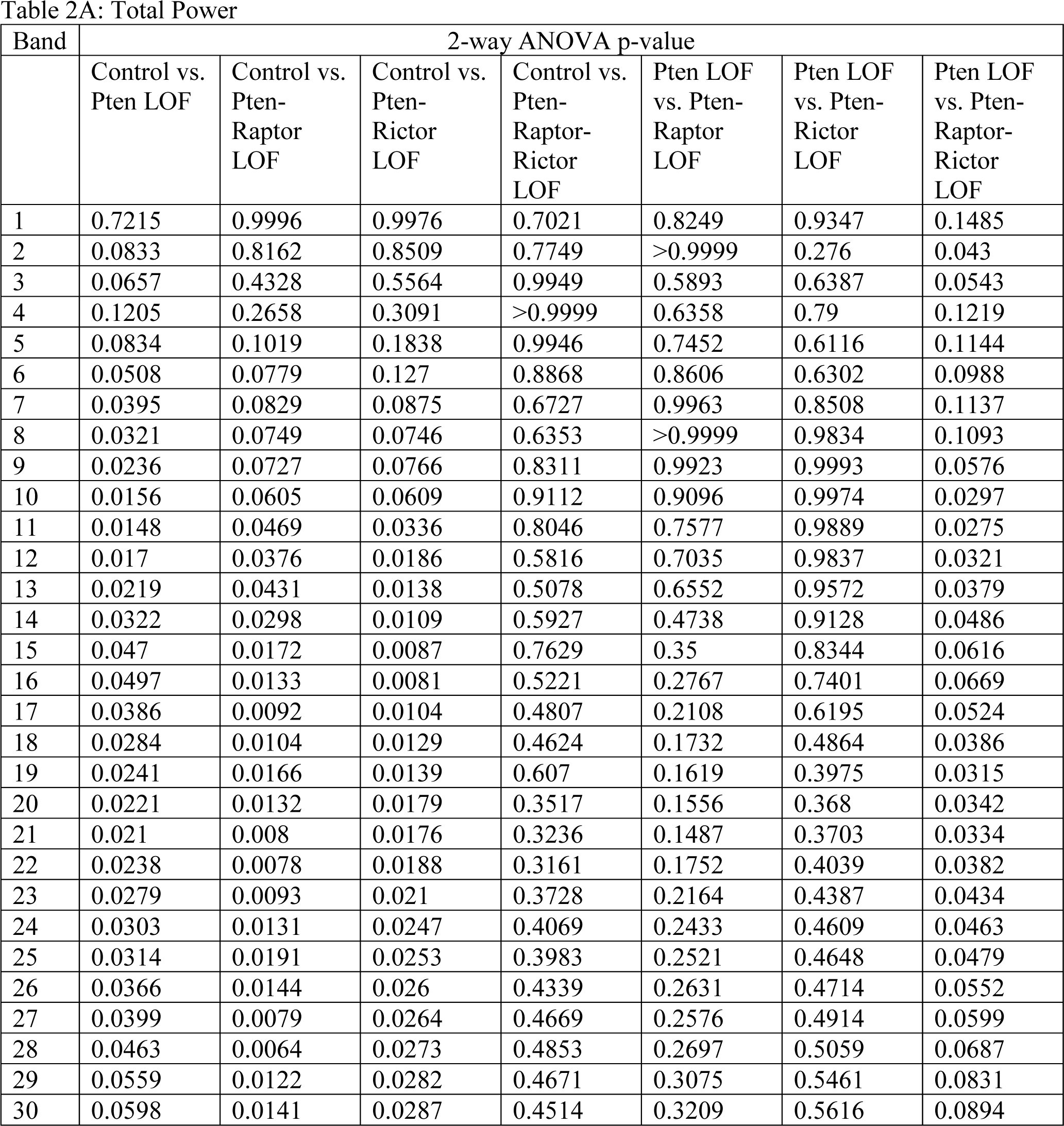
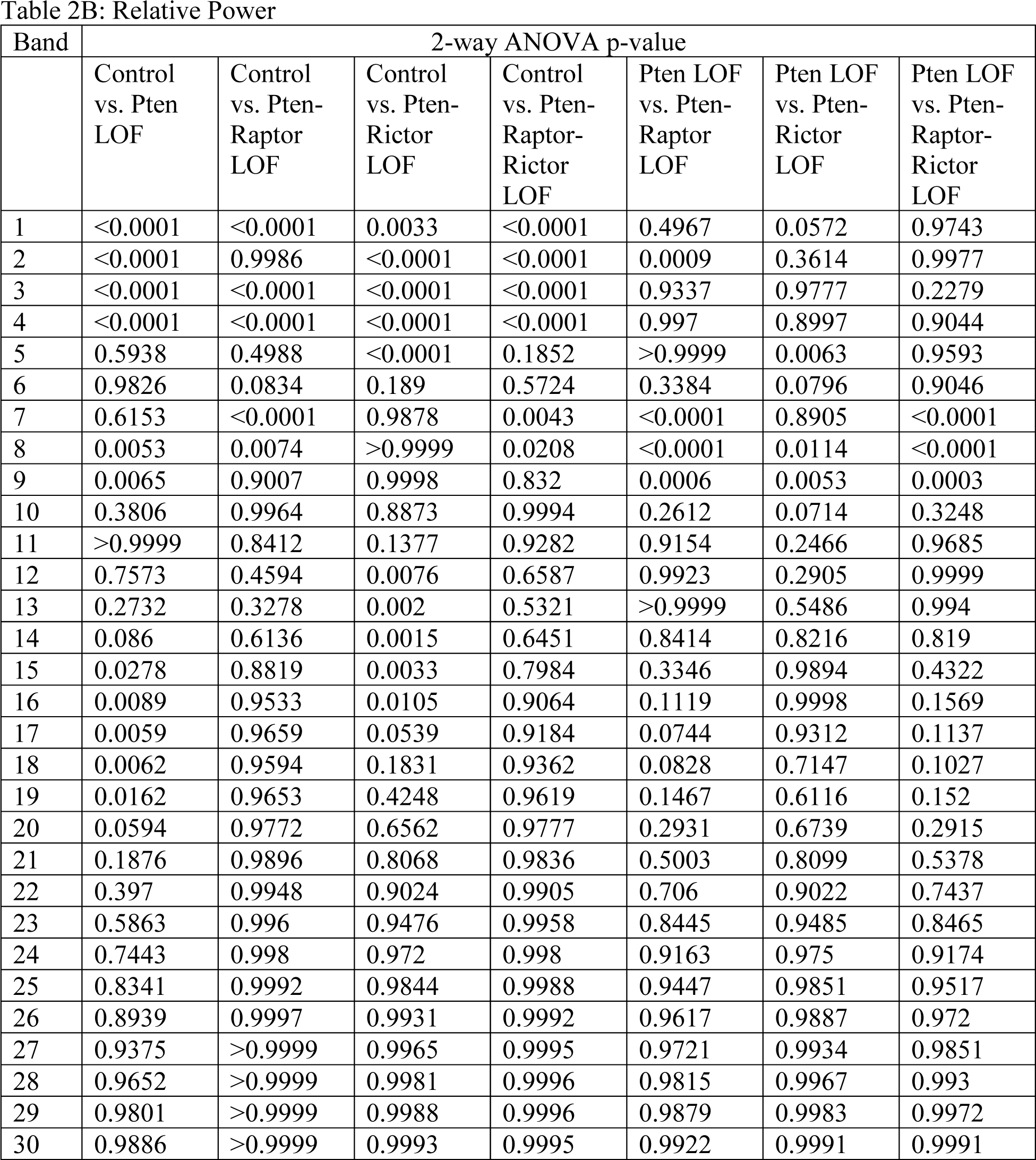
EEG power.

**Table 3.**
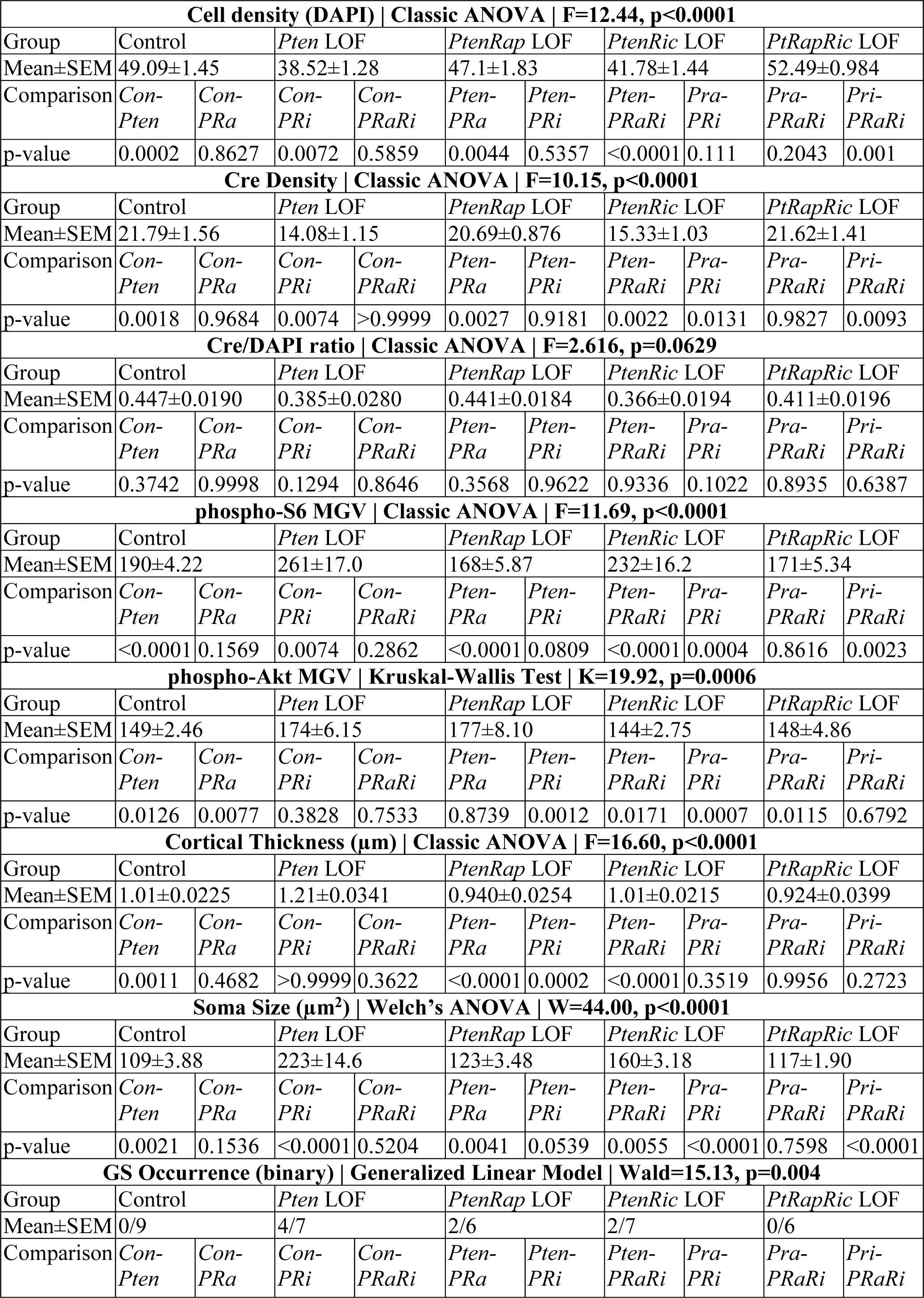
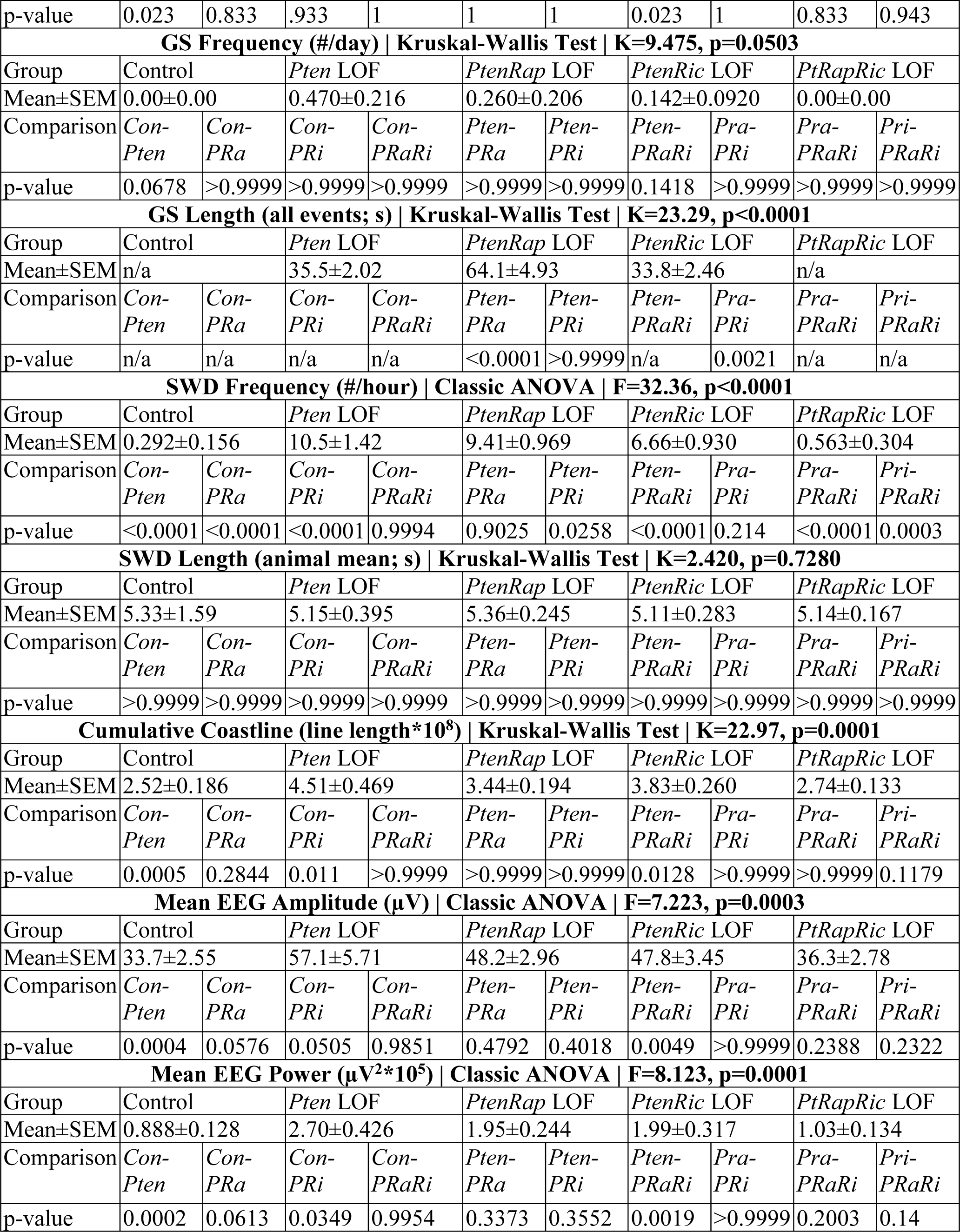
Results of Statistical Tests.

## Discussion

Convergent discoveries have positioned mTORopathies as prime candidates for a precision medicine approach [30,31]. There has been some clinical success with rapamycin analogues [32], but preclinical animal studies have suggested ways to further improve the selectivity and efficacy of the targets [12]. For the subset of mTORopathies in which mTORC1 but not mTORC2 signaling is hyperactive, selective inhibition of mTORC1 and its downstream effectors rescues many animal phenotypes, including seizures [33,34]. Studies using rapamycin as an mTORC1 inhibitor also came to this conclusion in *Pten* LOF models, but potential inhibition of mTORC2 and the finding that genetic inhibition of mTORC1 via *Rptor* deletion did not stop seizures, whereas inhibition of mTORC2 did [21], challenged this view. Here, we tested whether mTORC1 or mTORC2 inhibition alone were sufficient to block disease phenotypes in a model of somatic *Pten* LOF. Neither was, which agrees with previous findings that seizures persist after *Rptor* LOF and *Rictor* LOF in separate model systems [21,22]. This data could be interpreted as suggesting that epilepsy downstream of *Pten* LOF can proceed via mTORC-independent mechanisms, such as elevated ß-catenin or its protein phosphatase activity [25,35]. However, we found that simultaneous mTORC1 and mTORC2 inhibition corrected all of the phenotypes of our model except for a rightward shift in relative EEG power (Fig. 4B). This suggests that epilepsy can be caused by hyperactivity of either mTORC1 or mTORC2-dependent signaling downstream of *Pten* LOF, but not alternative pathways. Future studies should determine whether this is also true for other gene variants that hyperactivate both mTORC1 and mTORC2, as well in other models that have fully penetrant generalized seizures.

Epilepsy in Pten-Ric LOF mice likely emerges via mechanisms similar to those in mTOR pathway variants such as TSC that hyperactivate mTORC1 but not mTORC2, although additional effects of mTORC2 inactivation may contribute [22]. Epilepsy in Pten-Rap LOF occurs in the absence of macrocephaly, a hallmark of mTORC1 hyperactivity, possibly due to mTORC2 hyperactivity’s effect on synaptic transmission [20]. Roles for both complexes in emergent neural network function are corroborated by findings that behavior can be altered by either mTORC1 or mTORC2 inactivation [36]. Rapamycin derivatives are somewhat selective for mTORC1 [37,38] and are currently being tested in the clinic to treat mTORopathies and PTEN-related disorders [39]. Even more selective targeting of mTORC1 than can be achieved through rapamycin and its derivatives has been proposed as a way to effectively treat disease with fewer side effects [40]. Our data suggest that this may only be true for mTORopathies that selectively hyperactive mTORC1. For others, including *PTEN*, *PIK3CA*, and *AKT*, dual inhibitors or inhibitors of PI3K, which are being tested preclinically [41–43], are likely to be required.

## Materials and Methods

### Animals

*Rptor* (Jackson Labs #013188) and *Rictor* (Jackson Labs #020649) homozygous floxed mice were crossed to a *Pten* homozygous floxed line ([44] Jackson Labs #006440).

Experimental animals were the offspring of two animals homozygous floxed for *Pten* and heterozygous floxed for *Raptor* and/or *Rictor*. Intracranial viral injections were done at P0 or P1. Animals that were homozygous for the floxed *Pten* allele and either homozygous or wild-type at the *Raptor* and/or *Rictor* allele were injected with an AAV9 viral vector expressing an eGFP-Cre fusion driven by the *hSyn* promoter (Addgene #105540). Control animals were of the same genotype but were injected with an AAV9 virus expressing eGFP under control of the *hSyn* promoter, but not Cre (Addgene #105539). Pup injections were conducted with hypothermic anesthesia. A Nanoject (Drummond) was used to deliver 100 nL of Cre or control virus to three sites spanning the left hemisphere of the cortex. Pups were quickly rewarmed and returned to their home cage. Additional control animals with no floxed genes on a C57B6/J background (Jackson Labs #000644) were injected with the eGFP-Cre expressing AAV9 to ensure that Cre expression did not have effects independent of the *Pten^fl/fl^*genotype (Supplementary Figure 3).

### Surgeries and EEG Recordings

Animals were aged to six weeks and then implanted with a wireless EEG monitoring headcap. Animals were anesthetized with 4% isofluorane and arranged on a stereotaxic surgical apparatus with ear bars. Anesthesia was maintained with 1.5-2.5% isofluorane. The skull surface was exposed and holes were made with a 23g needle at six locations on the skull surface. 3/32” screws (Antrin Miniature Specialties) were inserted into the skull to record epidural EEG. Screws were secured with VetBond then covered with surgical cement. Recording leads were placed bilaterally at approximately Bregma AP +1.0 ML 1.0, AP -1.0 ML 1.5, and AP -2.5 ML 1.5. The screws were attached to a six-pin Millimax strip which was secured to the skull with additional cement. Animals were administered 5 mg/kg ketoprofen post-surgery and allowed to recover for at least 5 days before EEG recording. Surgical protocol was based on guidance provided by Pinnacle Technologies (https://www.pinnaclet.com).

After recovery from surgery, animals were fitted with wireless three-channel EEG preamplifiers (Pinnacle Technologies) and EEG signal was recorded at 256 Hz with simultaneous video recording. EEG recording session length was dependent on battery life. A total of at least 150 hours of EEG data was collected for each experimental animal (Table 1).

### Histology and imaging

EEG-implanted animals were euthanized by isofluorane overdose followed by transcardiac perfusion with 4% paraformaldehyde. Brains were postfixed in 4% paraformaldehyde for 24 hours and then transferred to 30% sucrose for at least 48 hours. 40 µm coronal slices were cut and preserved in a cryoprotectant solution (50% PBS, 30% ethylene glycol, 20% glycerol). Prior to staining, slices were washed three times with PBS. Eight unstained slices spanning the affected area (Bregma -3.6 through Bregma +1.1) were mounted with DAPI Fluoromount-G (SouthernBiotech) and used to assess cortical thickness, GFP expression, and cell density. GFP expression, cell density, and soma size were measured in the upper layers of the cortex to reduce within-subject variability. GFP expression was also measured in the dentate gyrus, CA1, and CA3 regions of the dorsal hippocampus. For other analyses, three sections per animal at approximately Bregma -1.6, -2,1, and -2.6 were used. A fluorescent Nissl stain (Invitrogen N21482, 1:50) was used to assess soma size.

After washing, slices were placed in blocking solution (10% normal goat serum, 0.1% Triton X-100, and PBS) for 1 hour. Slices were incubated in the following primary antibodies for 3 hours at room temperature: phospho-S6 Ribosomal Protein Ser240/244 (rabbit monoclonal, 1:1000 dilution, Cell Signaling Technology, catalog #5364, RRID: AB_10694233), phospho-AKT Ser473 (rabbit monoclonal, 1:1000 dilution, Cell Signaling Technology, catalog #4060), and NeuN (guinea pig polyclonal, 1:1000 dilution, Synaptic Systems, catalog #266004). Following primary antibody application, slices were washed three times in PBS and the incubated in AlexaFluor secondary antibodies goat anti-guinea pig 647 and goat anti-rabbit 594 (Invitrogen) for 1 hour at room temperature.

Slides stained for Nissl substance, NeuN/pS6, and NeuN/pAkt were imaged at 10x (2048x2048) with 5 µm z-step on a Nikon C2 confocal microscope (UVM Microscopy Imaging Core). One image of the left (GFP-expressing) cortical hemisphere in each of three region-matched slices per animal was collected for analysis. Widefield images (2560x2160) of unstained DAPI-mounted cross-sectional slices were taken with a Zyla sCMOS camera (Andor) mounted on an upright microscope (IX73; Olympus) at 5x and 20x resolution. Pixel width was manually calculated using an 0.1 mm hemocytometer.

### Slice electrophysiology

Slice electrophysiology was conducted at P14-30. Animals were deeply anesthetized and decapitated, and the brain was quickly dissected into ice-cold cutting solution (126 mM NaCl, 25 mM NaHCO3, 10 mM d-glucose, 3.5 mM KCl,1.5 mM NaH2PO4, 0.5 mM CaCl2, 10.0 mM MgCl2, pH 7.3-7.4). 350-um slices were cut using a Leica 1000S Vibratome. Slices were transferred to 37°C aCSF (126 mM NaCl, 3.5 mM KCl, 1.0 mM MgCl2, 2.0 mM CaCl2, 1.5 mM NaH2PO4,25 mM NaHCO3, and 10 mM d-glucose, pH 7.3-7.4) and incubated for 30 min. The slices were then incubated at room temperature for at least another 30 min prior to recording. All solutions were continuously bubbled with 95% O2 and 5% CO2. Individual slices were transferred to a recording chamber located on an upright microscope (BX51; Olympus) and were perfused with heated (32-34°C), oxygenated aCSF (2 mL/min).

Whole-cell voltage-clamp and current-clamp recordings were obtained using Multiclamp 700B and Clampex 10.7 software (Molecular Devices). GFP+ cells in cortical layer 2/3 in the motor cortex were targeted for patching. Intracellular solution contained (in mM): 136 mM K-gluconate, 17.8 mM HEPES, 1 mM EGTA, 0.6 mM MgCl2, 4 mM ATP, 0.3 mM GTP, 12 mM creatine phosphate, and 50 U/ml phosphocreatine kinase, pH 7.2. When patch electrodes were filled with intracellular solution, their resistance ranged from 4–6 MΩ. Access resistance was monitored continuously for each cell.

For current-clamp experiments, the intrinsic electrophysiological properties of neurons were tested by injecting 500-ms square current pulses incrementing in 20 pA steps, starting with -100 pA. The membrane time constant was calculated from an exponential fit of current stimulus offset. Input resistance was calculated from the steady state of the voltage responses to the hyperpolarizing current steps. Membrane capacitance was calculated by dividing the time constant by the input resistance.

Action potentials (APs) were evoked with 0.5 s, 20 pA depolarizing current steps. Rheobase was defined as the minimum current required to evoke an AP during the 500 ms of sustained somatic current injection. For voltage-clamp experiments to measure spontaneous EPSC frequency and amplitude, neurons were held at -70 mV and recorded for 2 min. All electrophysiology data were analyzed offline with AxoGraph X software (AxoGraph Scientific).

### EEG analysis

EEG files were converted from Pinnacle’s proprietary format to European Data Format (EDF) files and imported to Matlab. A filtering program was used to flag traces in which seizures were likely to be occurring based on amplitude and line length between data points. For Control and PtRapRic LOF animals, all 10 second epochs in which signal amplitude exceeded 400 µV at two time points at least 1 second apart were manually reviewed by a rater blinded to genotype. This method was not suitable for the *Pten* LOF, *Pten-Raptor* LOF, and *Pten-Rictor* LOF animals because of the density of high-amplitude interictal activity they displayed. For these animals, at least 100 of the highest-amplitude traces were manually reviewed and then traces with persistent abnormally low amplitude, often indicating postictal suppression, were reviewed as well. Flagged traces were displayed for a rater to mark the beginning and end of each seizure. We also reviewed at least 48 hours of data from each animal manually. All seizures that were identified during manual review were also identified by the automated detection program. Spike-and-wave discharge events were manually marked in 8 evenly spaced one-hour epochs in each of two 24-hour recording sessions per animal and verified on video not to be caused by movement such as chewing or scratching.

Baseline EEG measures were taken from a representative sample of EEG files for each animal.

692 evenly spaced five-second epochs were sampled over 24 hours, repeated for two recording sessions per animal. Line length was defined as the sum of the linear distances between adjacent data points during the five-second analysis epoch. Mean amplitude was defined as the mean of the absolute value of data points in the five-second analysis epoch. Power spectral density was calculated with the pwelch function in Matlab. All three EEG channels were analyzed and the mean of all channels was used for statistical analysis.

### Image Analysis

Image analysis was conducted using ImageJ/Fiji. Cell density and Cre expression in the cortex were automatically assessed using the Fiji Analyze Particles tool. Cre expression in the hippocampus was manually assessed by counting Cre-expressing punctae within a 100 µm linear portion of the region of interest. Because neuronal somas within the hippocampal cell layers are so closely packed together, we were unable to resolve DAPI punctae to assess cell density in the hippocampus. Soma size was measured by dividing Nissl stain images into a 10 mm^2^ grid. The somas of all GFP-expressing cells fully within three randomly selected grid squares in Layer II/III were manually traced. pS6 and pAkt expression were measured by drawing 200 µm wide columns spanning all cortical layers. Background was subtracted from images with a 30 µm rolling ball algorithm (Fiji/ImageJ). The mean pixel value of the column was recorded and values were averaged by animal.

### Statistical analysis

Prism 9 or 10 (GraphPad Prism) was used to conduct statistical analyses and create graphs. All data are presented as mean ± SEM. Power analyses on preliminary and published data were used to calculate the number of animals necessary for this study, with the exception of GS occurrence, which is discussed below. All data distributions were assessed for normality using the Shapiro–Wilk test. If data did not meet the criteria for normal distribution (p<0.05), Kruskal-Wallis tests were used to assess statistical relationships with Dunn’s post-hoc correction for multiple comparisons. Variance was assessed using Brown-Forsythe tests. If variance differed significantly (p<0.05), Welch’s ANOVA test was used to assess statistical relationships.

Group differences in normally distributed datasets with equal variances were assessed using one-way ANOVA with Tukey post-hoc correction for multiple comparisons. Power spectral density was assessed using two-way ANOVA with Tukey post-hoc analysis. Seizure occurrence was assessed with a Generalized Linear Model with a binary distribution and logistic link function implemented in SPSS and Bonferroni corrected for multiple comparisons. See Tables 2 and 3 for details regarding statistical tests. A power analysis showed that an excessive number of animals would be required to detect a decrease in GS incidence or frequency by *Rptor* or *Rictor* LOF in this model, thus we chose not to pursue the possibility that *Rptor* or *Rictor* loss decreases these outcome measures.

## Acknowledgements

Research reported in this publication was supported by NINDS grant R01NS110945 (M.C.W.), as well as P20GM135007, Core C: Customized Physiology and Imaging Core. Images created with BioRender. We thank Caitlynn Barrows, Elise Prehoda, and Willie Tobin for early experiments helping create the mouse model.

---

**Supplementary Figure 1.**
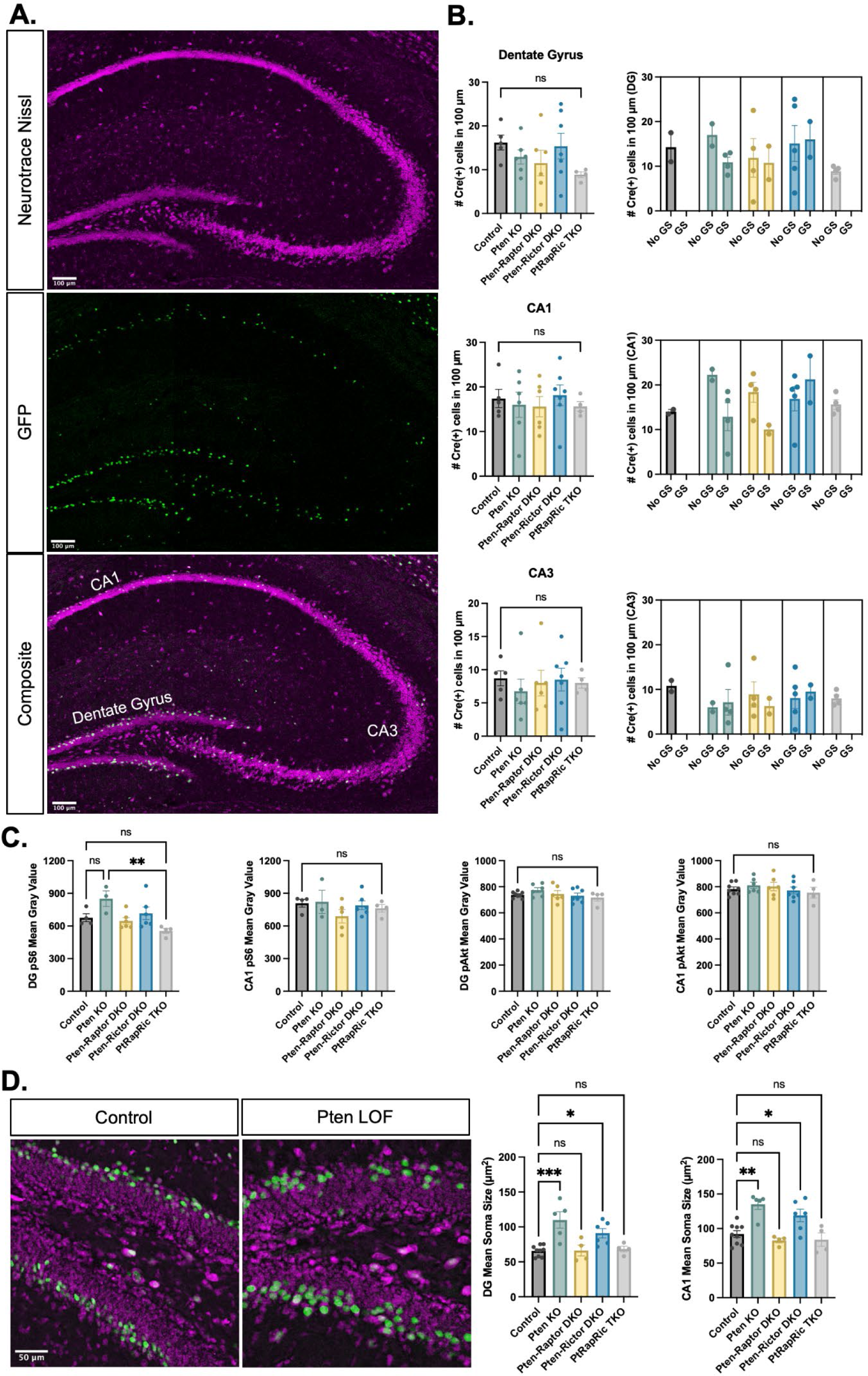
Analysis of Cre expression in the hippocampus and its impact on outcome measures. Although the virus injection was targeted to the cortex, sparse GFP expression was also observed in the dorsal hippocampus, most prominently in CA1 and the dentate gyrus. A) Representative images showing GFP expression in the dorsal hippocampus. B) Summary data showing that hippocampal Cre expression did not differ between groups and was not related to generalized seizure development. C) phospho-S6 and phospho-Akt in the hippocampus were not significantly elevated from Control values in any group, but PtRapRic LOF animals had significantly lower phospho-S6 expression than Pten LOF animals. D) Representative images and summary data showing that soma size in the hippocampus was increased in Pten LOF and PtenRic LOF animals, as was also observed in the cortex. Error bars show mean ± s.e.m. ns indicates p>0.05, * indicates p<0.05, ** indicates p<0.01, *** indicates p<0.001 as assessed by one-way ANOVA with Tukey multiple comparisons correction.

**Supplementary Figure 2.**
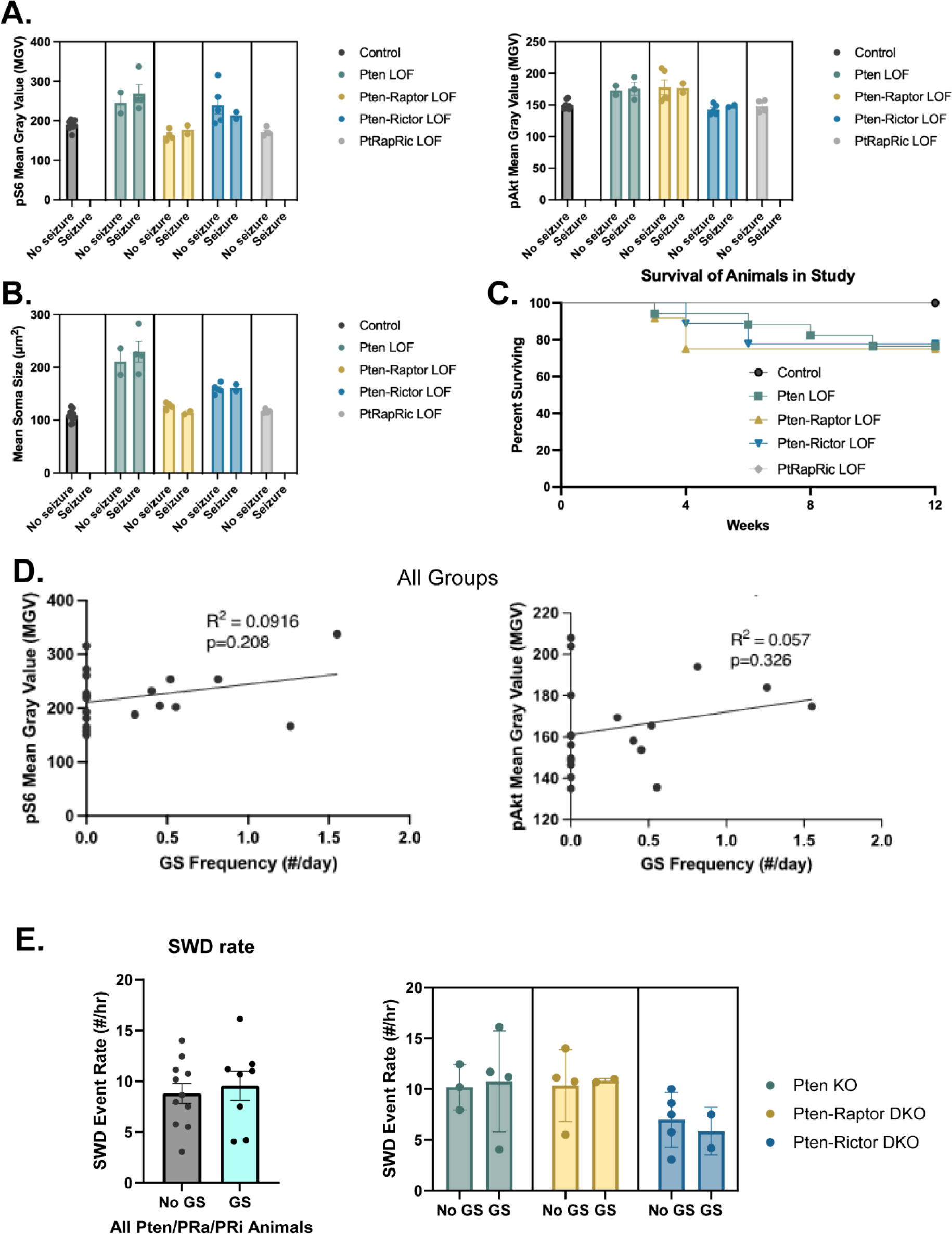
Focal cortical Pten LOF, Pten-Rap LOF, and Pten-Ric LOF cause a spectrum of outcomes. A subset of animals in the Pten LOF, Pten-Rap LOF, and Pten-Ric LOF groups displayed spontaneous epileptiform activity, but not generalized seizures. When compared side by side, animals within each genotype that did and did not display generalized seizures showed similar mTOR pathway activity levels (A) and soma sizes (B). Survival plot (C) shows survival of animals in the study by genotype. Some animals in the Pten LOF, Pten-Rap LOF, and Pten-Ric LOF groups were found dead during the study, but no deaths were observed in Control or PtRapRic LOF groups. Mortality often occurred prior to EEG recordings, so we could not ascertain whether early mortality was associated with generalized seizures. When all groups with GS were plotted together, there was no significant correlation between pS6 or pAkt levels and the presence of GS (D). There was also no apparent relationship between GS presence and SWD rate (E).

**Supplementary Figure 3.**
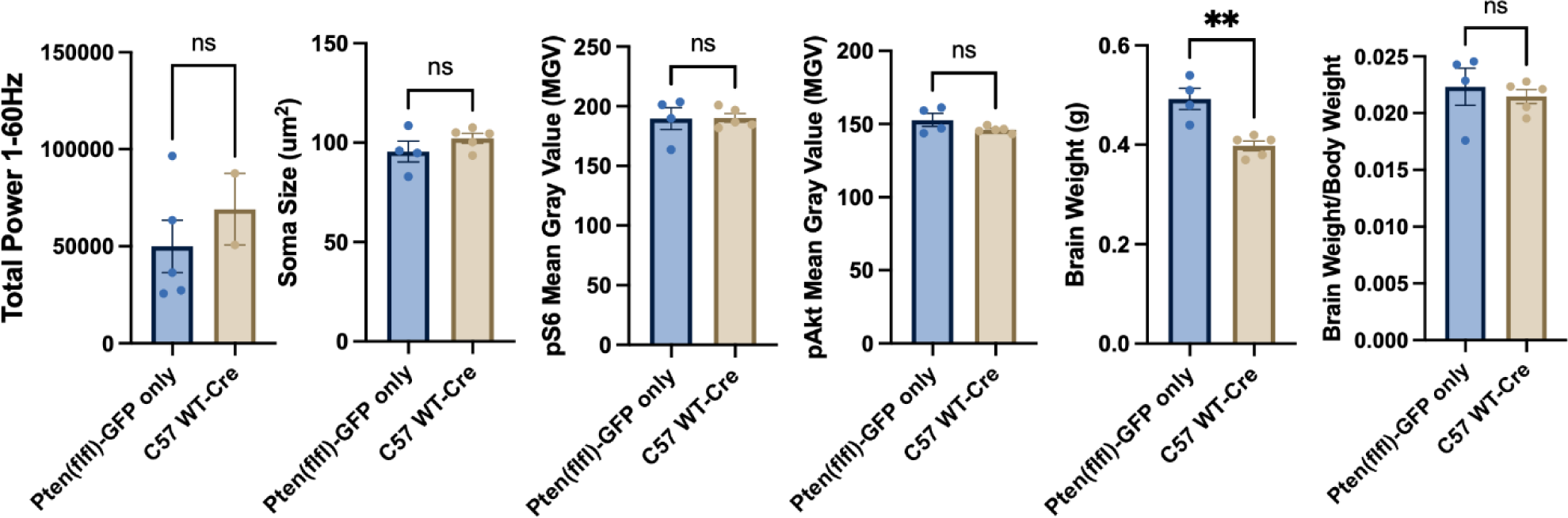
Cre virus exposure does not significantly impact cortical morphology or baseline EEG. C57B6/J mice with no floxed genes were injected with the Cre virus as an additional control group. These mice did not have GS or SWDs. They were also not different from *Pten*^fl/fl^ injected with the Control virus, except for significantly different brain weights. The brain weight/body weight ratio did not differ between the groups.

